# GP-ML-DC: An Ensemble Machine Learning-Based Genomic Prediction Approach with Automated Two-Phase Dimensionality Reduction via Divide-and-Conquer Techniques

**DOI:** 10.1101/2024.12.26.630443

**Authors:** Quanzhong Liu, Haofeng Ma, Zhuangbiao Zhang, Zhunhao Hu, Xihong Wang, Ran Li, Yudong Cai, Yu Jiang

**Affiliations:** College of Information Engineering, Northwest A&F University, Yangling, 712100, China; Key Laboratory of Animal Genetics, Breeding and Reproduction of Shaanxi Province, College of Animal Science and Technology, Northwest A&F University, Yangling 712100, China; Key Laboratory of Livestock Biology, Northwest A&F University, Yangling, Shaanxi 712100, China; Shaanxi Engineering Research Center of Agricultural Information Intelligent Perception and Analysis, Northwest A&F University, Yangling 712100, China

## Abstract

Traditional machine learning (ML) and deep learning (DL) methods for genome prediction often face challenges due to the imbalance between the limited number of samples (*n*) and the large number of single nucleotide polymorphisms (SNPs) (*p*), where *n* is much smaller than *p*. To address this, we propose GP-ML-DC, an innovative genome predictor that combines traditional ML and DL models with a unique two-phase, parameter-free dimensionality reduction technique. Initially, GP-ML-DC reduces feature dimensionality by characterizing genes as features. Building on big data methodologies, it employs a divide-and-conquer approach to segment gene regions into multiple haplotypes, further decreasing dimensionality. Each haplotype segment is processed by a sub-task based on traditional ML, followed by integration via a neural network that synthesizes the results of all sub-tasks. Our experiments, conducted on four cattle milk-related traits using ten-fold cross-validation and independent testing, show that GP-ML-DC significantly surpasses current state-of-the-art genome predictors in prediction performance.

## Introduction

Predicting phenotypes directly from genotypes offers significant advantages in modern breeding [1]. To address the limitations of traditional marker-assisted selection (MAS) for predicting complex traits in crop and livestock breeding, genomic selection (GS) [2] has been introduced. GS utilizes dense molecular markers across the entire genome to evaluate the breeding values of individuals. The initial step in GS involves building a genomic prediction (GP) model using the training population, thereby establishing genotype-phenotype relationships. Once constructed, this model can predict phenotypes or breeding values for test populations based solely on their genotypic data. Commonly, GP models employ statistical methods, such as linear mixed models [2], including best linear unbiased prediction (BLUP), genomic BLUP (GBLUP), HIBLUP [3], and Bayesian linear regression models [4], which have been extensively applied in both animal and crop breeding [5, 6].

Linear mixed models have indeed propelled advancements in breeding by capturing linear relationships; however, they often fail to account for non-linear effects on complex traits [7]. Recently, machine learning (ML)-based GP methods have emerged, enhancing predictions of complex traits due to their ability to capture non-linear relationships. Performance comparisons among traditional linear models, conventional ML, and deep learning (DL)-based models indicate that no single approach consistently outperforms others universally [8]. Reviews [1, 5, 7, 9, 10] of diverse ML and DL predictors for GP similarly conclude that no single method excels across all contexts, underscoring the ongoing challenge of predicting complex phenotypes.

Despite advancements, ML and DL methods face challenges due to the imbalance where the number of individuals (*n*) is significantly less than the number of SNPs (*p*) [5, 7], hindering the performance of both statistical and ML models [11]. To address this, feature selection algorithms are often employed to reduce SNP dimensionality and improve GP performance [12]. Yet, standard feature selection algorithms such as filter, wrapper, and embedded methods [13] face difficulties in fully enhancing performance and often overlook non-linear SNP relationships or depend on a threshold of dimensionality reduction. While haplotype-based methods reduce dimensionality effectively [14, 15], they depend on hyperparameters for defining haplotype block boundaries such as window size or an LD threshold [16]. Using genes as feature descriptors can somewhat reduce dimensionality [17, 18]; however, the dataset’s dimensionality, indicated by the number of annotated genes (*q*), often remains high (*n* ≪ *q*). Consequently, the issue of having more features than samples (*n* ≪ *q*) persists. Addressing these high-dimensional features continues to pose a significant challenge for ML-based GP.

To resolve these issues, we propose a machine learning-based genomic predictor, GP-ML-DC, aimed at enhancing GS performance. GP-ML-DC introduces a parameter-free, gene-based feature selection algorithm, reducing dimensionality from SNPs to genes. Inspired by big data principles, it employs a divide-and-conquer strategy [19] to split gene regions into 16 core haplotypes, treating predictions from each as meta-features. This results in two phases of automatic dimension reduction: from *p* (the number of SNPs) to *q* (the number of genes), and from *q* to 16 meta-features, which are then inputted into a neural network for final phenotype predictions. GP-ML-DC is designed for end-to-end application. We conducted a five-time ten-fold cross validation on four milk-related traits (DMY, MFY, MPY, and SCS) using datasets from dairy farms in Hebei province, China, and compared GP-ML-DC with three state-of-the-art predictors including GBLUP (based on statistics models), LightGBM [20] (based on traditional ML), and DNNGP [21] (based on DL). Results demonstrate that GP-ML-DC improved predictions by 15.2% for DMY, 21.4% for MFY, 10.0% for MPY, and 19.5% for SCS compared to LightGBM (the second-best predictor). Independent test using datasets from dairy farms in Ningxia province, China indicate that GP-ML-DC improved predictions by 24.2% for DMY, 19.1% for MFY, 17.9% for MPY, and 8.4% for SCS compared to LightGBM (the second-best predictor).

## Results

### Design of genome prediction model

**Figure 1** outlines the structure of the proposed GP-ML-DC model, which consists of two main modules: data mapping and ensemble ML-based prediction. The data mapping module involves two stages: feature engineering and data division. In the data mapping phase, we first perform feature engineering by extracting 16 features for each gene from variant information files. This yields a matrix *M* of size *q* × 16 for each sample, where *q* denotes the number of genes. The raw data from *n* samples is thereby transformed into a three-dimensional data tensor *D*(1.. *n*, 1.. *q*, 1. .16), thereby achieving dimensionality reduction from *p* to *q* (called the first phase of dimensionality reduction). We then perform the data division by extracting a matrix *D*(:, :, *f*) for the *f*th feature from *D*, with *f* ∈ [1,16].

**Figure 1.**
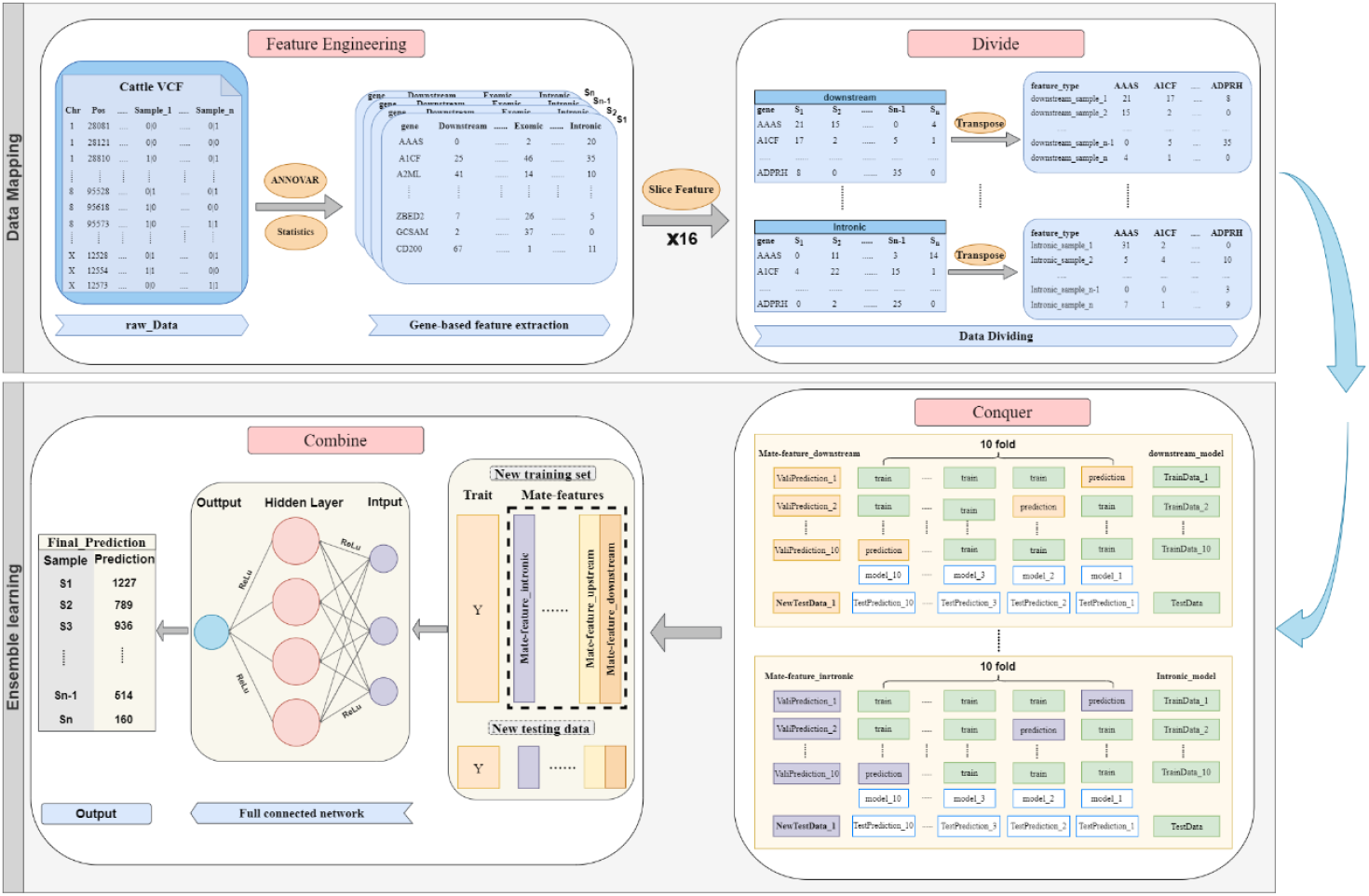
The workflow of GP-ML-DC

In the ensemble ML-based prediction module, there are two stages: conquering and combining. During the conquering stage, a LASSO model is trained on the extracted *D*(:, :, *f*) for each feature *f* to learn a meta-feature using a cross-validation strategy from stacking ensemble learning [22]. The 16 meta-features are then combined to construct a training set for the combining stage, further reducing feature dimensionality from *q* to 16 (called the second phase of dimensionality reduction). In the combining stage, a densely connected network is subsequently trained on this dataset to predict testing samples. Further details can be found in the **Methods** section.

### Performance comparison with other methods across ten-fold cross validation

We conducted five rounds of ten-fold cross-validation on four traits (DMY, MFY, MPY, and SCC) using cow datasets (refer to **Methods**) from Hebei province, China. The performance of GP-ML-DC was evaluated against three state-of-the-art methods: the GBLUP statistics model, LightGBM [20] based on traditional ML, and DNNGP [21] based on DL. As shown in **Figure 2A**, GP-ML-DC outperformed GBLUP (0.598 versus 0.499 on DMY, 0.569 versus 0.419 on MFY, 0.570 versus 0.477 on MPY, 0.388 versus 0.342 on SCS), LightGBM (0.598 versus 0.576 on DMY, 0.569 versus 0.545 on MFY, 0.570 versus 0.567 on MPY), and DNNGP (0.598 versus 0.494 on DMY, 0.569 versus 0.447 on MFY, 0.570 versus 0.438 on MPY, 0.388 versus 0.373 on SCS) in terms of PCC (Pearson’s correlation coefficient). Additionally, as shown in **Figure 2B**, GP-ML-DC also surpassed LightGBM (the second-best performer) concerning R^2^ values (0.358 versus 0.332 on DMY, 0.324 versus 0.299 on MFY, 0.328 versus 0.323 on MPY).

**Figure 2.**
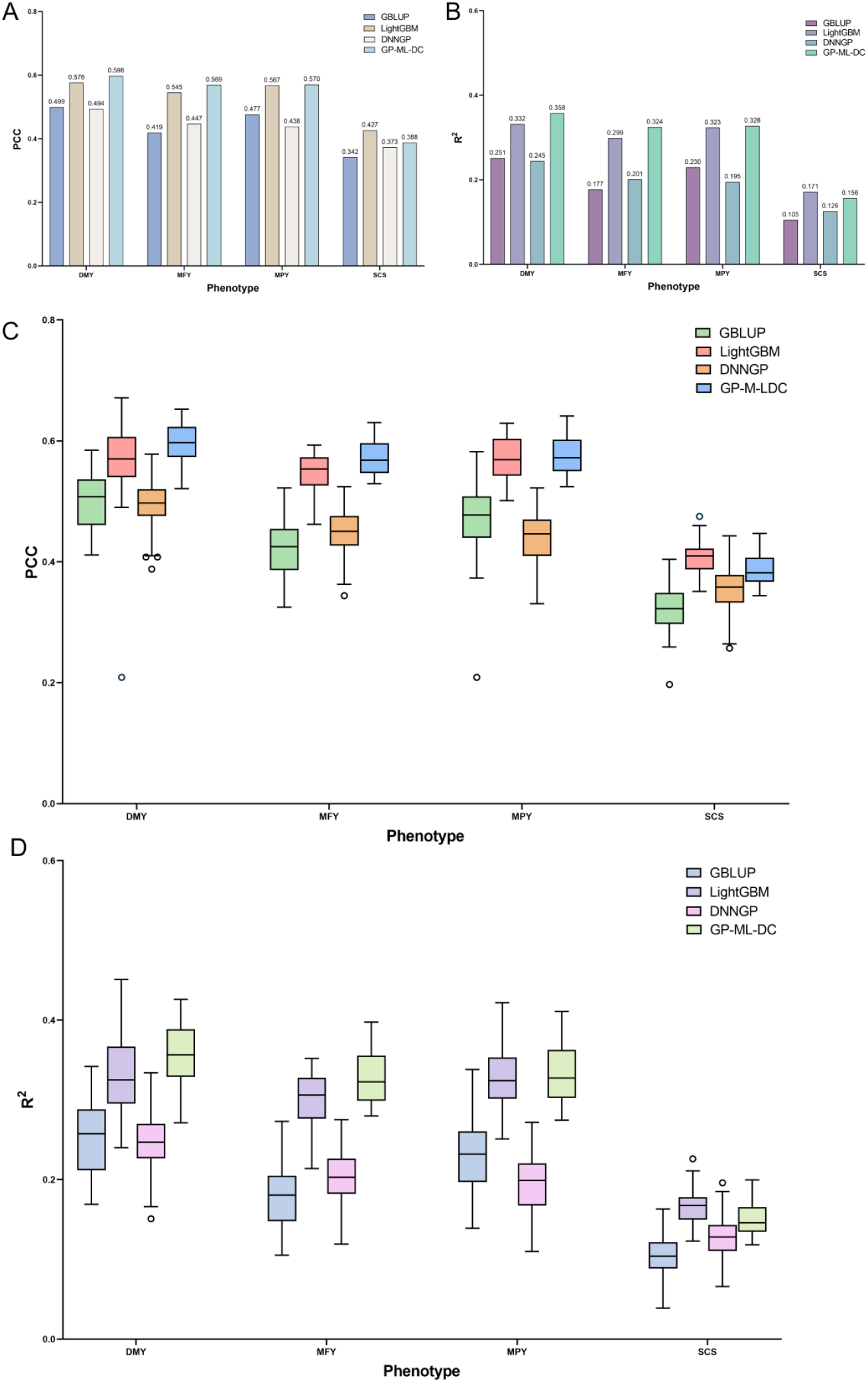
Performance comparison of four methods across four traits from the Hebei dataset using five-time ten-fold cross-validation. (A) Comparation results for the average PCC. (B) Comparation results for the average R^2^. (C) Distribution of 50 PCC scores from five-time ten-fold cross-validation. (D) Distribution of 50 R^2^ scores from five-time ten-fold cross-validation.

To further validate the model’s robustness, we plotted 50 prediction results from five rounds of ten-fold cross-validation, as shown in **Figure 2C** and **Figure 2D**, for PCC and R^2^ performance, respectively. As can be seen, compared with other three methods, GP-ML-DC demonstrated higher median PCC and *R*^2^ values across 50 iterations for DMY, MFY, and MPY. Additionally, we further conducted a Levene’s test and Student’s t-test to examine the statistical differences in performance between GP-ML-DC and other three predictors, with a significance cut-off of 0.05. Statistical results for PCC performance are provided in **Tables S1-S3**, and for *R*^2^ performance are provided in **Table S4-S6**. In terms of PCC, variance differences were significant across all four traits between GP-ML-DC and GBLUP (**Table S1**), as well as three traits (DMY, MFY, and MPY) between GP-ML-DC and LightGBM (**Table S2**). In contrast, mean differences were statistically significant across four traits in **Table 1**, and for DMY and MFY in **Table S2**. For the statistical results between GP-ML-DC and DNNGP (**Table S3**), variance differences were significant across the four traits; mean differences were statistically significant for each trait. Similar *R*^2^ results are observed in **Table S4-S6**, except for the SCS trait. These results indicated significant differences in the means of PCC and R^2^ in most traits between GP-ML-DC and the other methods, underscoring its more robust performance of GP-ML-DC.

### Independent testing and generalization performance comparison

Given the importance of generalization in ML models, we assessed how well GP-ML-DC transferred GP capabilities to different populations. Unlike traditional ML models, which rely on SNP-based matrices that may vary between groups, GP-ML-DC uses gene-based predictions that are applicable across diverse environments, such as populations from various farms. To assess the model’s generalization capability, we utilized a model trained on the Hebei dataset to independently predict four milk-related traits in a Ningxia dataset (refer to the **Methods** section). We compared the performances of GP-ML-DC with GBLUP, LightGBM, and DNNGP based on PCC and R^2^ values by comparing predicted and observed values. As illustrated in **Figure 3**, GP-ML-DC consistently outperformed other three methods in PCC and R^2^ values for MFY, and MPY traits. Specifically, GP-ML-DC showed superior performance compared to GBLUP, with PCC values of 0.198 versus 0.180 for MFY, and 0.229 versus 0.199 for MPY, as well as R^2^ values of 0.039 versus 0.034 for MFY, and 0.052 versus 0.040 for MPY. These findings indicate GP-ML-DC can effectively transfer GP across different populations on various farms.

**Figure 3.**
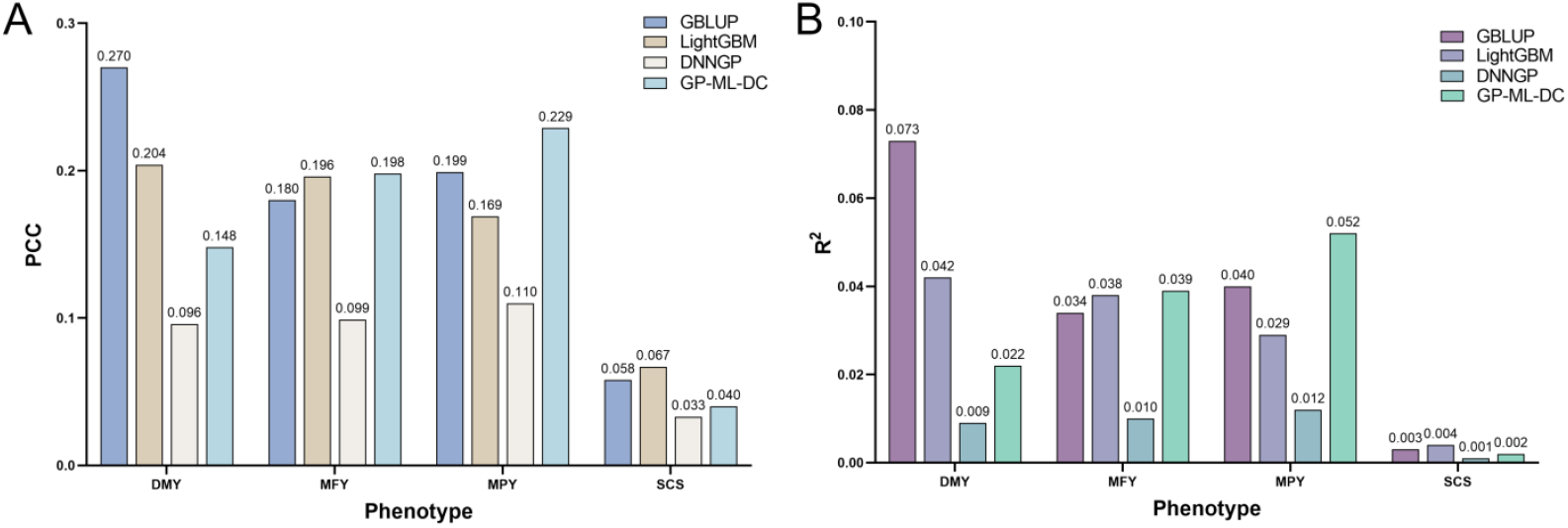
Performance comparison of four methods across four traits for the independent test on the Hebei dataset. (A) Comparation results for the PCC of four methods. (B) Comparation results for the R^2^ of four methods.

### Performance evaluation on the 50K SNP chip extracted by our model

We applied **Algorithm 4**, detailed in the **Methods** section, to the Hebei dataset to extract a 50K SNP chip, named SNP50N. To evaluate its effectiveness, we compared SNP50N with SNP50V3 [23], a widely recognized standard 50K SNP chip in the research community, using the same dataset. We performed ten-fold cross-validation tests using GBLUP on both SNP50N and SNP50V3 of the Hebei dataset. Results of these comparisons are presented are illustrated in **Figure 4**. As depicted, the GBLUP performance for SNP50N showed marked improvements in both PCC and R^2^ over SNP50V3 across all traits analyzed. For the DMY trait, enhancements were 10.3% and 21.4%, respectively (**Figure 4A-B**); for MFY, 11.8% and 24.3% (**Figure 4C-D**); for MPY, 10.6% and 22.2% (**Figure 4E-F**); and for SCS, 1.9% and 2.9% (**Figure 4G-H**).

**Figure 4.**
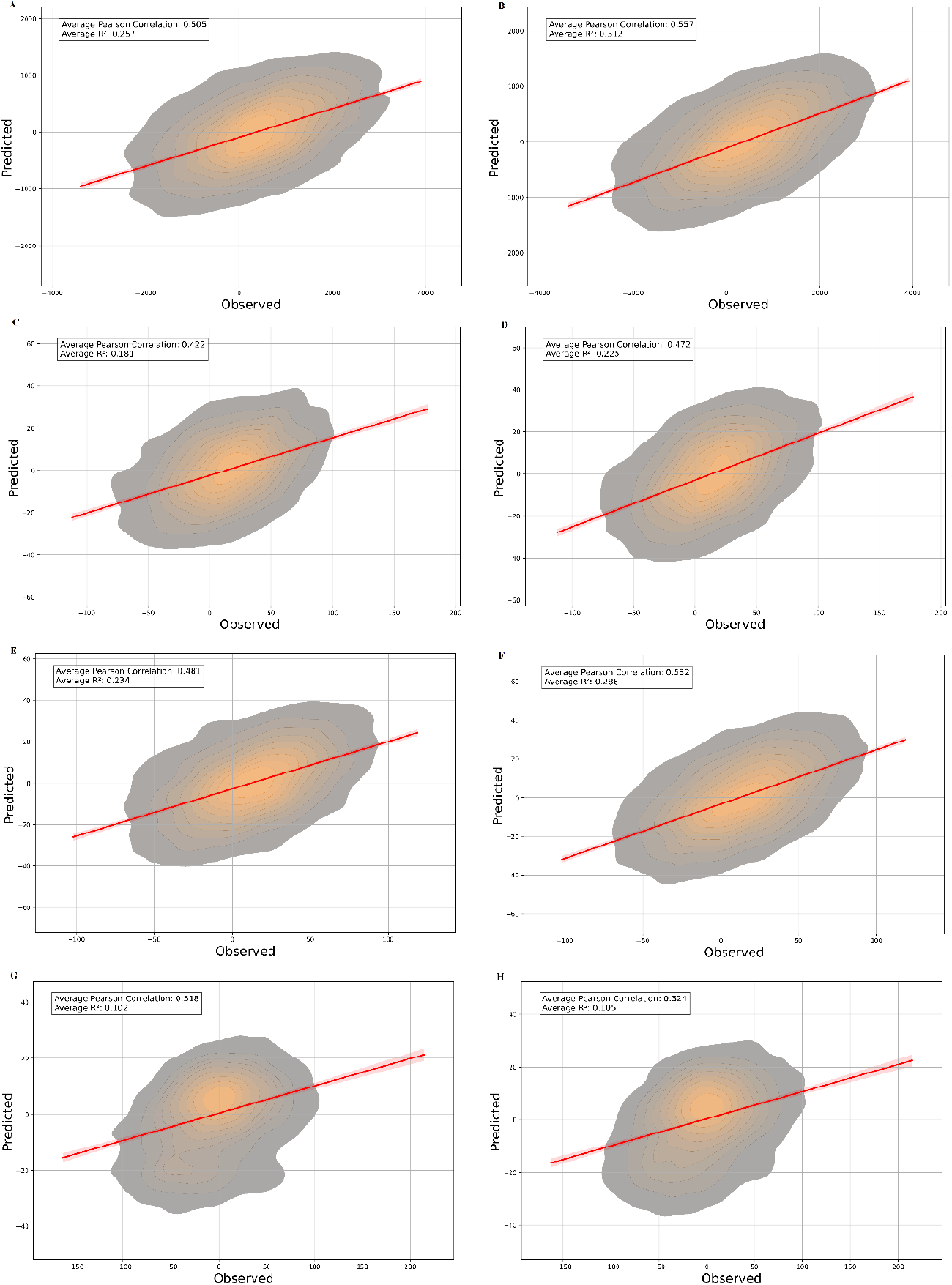
Performance comparisons of GBLUP between SNP50V3 and SNP50N extracted in this study. (A) Performances on SNP50V3 for the DMY trait. (B) Performances on SNP50N for the DMY trait. (C) Performances on SNP50V3 for the MFY trait. (D) Performances on SNP50N for the MFY trait. (E) Performances on SNP50V3 for the MPY trait. (F) Performances on SNP50N for the MPY trait. (G) Performances on SNP50V3 for the CSC trait. (H) Performances on SNP50N for the CSC trait.

## Discussion and conclusions

Genomic predictors often encounter challenges due to a limited number of samples compared to the vast number of genetic variants. Machine learning-based predictors also struggle with inconsistent SNP profiles across individuals or groups within the same species, limiting their broader applicability. Traditional feature selection methods attempt to address these issues by reducing the dimensionality of variables, but often at the expense of losing critical information necessary for effective genomic selection. In this study, we introduce GP-ML-DC, an innovative genome prediction model designed to address these challenges. This model embeds a novel two-phase, parameter-free dimensionality reduction process that utilizes genes to describe the genome, thereby enhancing its applicability across different individuals or groups within the species. Based on the concept of big data techniques, the approach goes beyond the traditional counting of SNPs in a gene region. It segments genes into haplotypes based on variant types using a divide-and-conquer strategy. Traditional machine learning models analyze each haplotype, while deep learning integrates these analyses to assess the collective contribution of all haplotypes to trait prediction. This ensemble strategy leverages the strengths of both traditional machine learning and deep learning in genomic selection.

Our experimental analysis, conducted through five rounds of ten-fold cross-validation, indicates that GP-ML-DC outperforms three other representative predictors, including those based on statistical, machine learning, and deep learning methods, for three out of four milk-related traits in experimental cow datasets. The 50K SNP chip produced by our method exhibits superior performance compared to the standard 50K SNP chip widely recognized in the research community. Additionally, the GP-ML-DC model can be seamlessly applied across diverse populations within the same species. Independent testing on milk-related trait prediction across various farms revealed that GP-ML-DC achieves better results compared to other predictors on two of four traits. A comparison between ten-fold cross-validation and independent testing reveals that GWAS (Genome-Wide Association Studies)-related SNP variants significantly influence model performance. This effect arises because these variants exhibit greater variability across distinct populations than across different folds within the same population. To our knowledge, GP-ML-DC is the first genome prediction tool that combines the concept of big data with artificial intelligence techniques. We anticipate that GP-ML-DC will establish itself as a leading genome prediction tool, providing accurate phenotype predictions and enhancing the breeding process.

## Methods

### Datasets

In this study, we collected data through the Dairy Herd Improvement (DHI) program from over 55,871 Holstein cows across Hebei and Ningxia provinces in China. Specifically, 26068 cows were sourced from nine large-scale farms in Ningxia, while 29083 cows came from four large-scale farms in Hebei. Our genetic evaluation focused on key milk production traits, including daily milk yield (DMY), milk fat yield (MFY), milk protein yield (MPY), and somatic cell score (SCS), utilizing a random regression test-day model [24]. From the overall dataset, we identified a subset of 6,411 cows possessing comprehensive pedigree and phenotypic records. Among these, 6,010 cows had documented first-lactation data for DMY, MFY, MPY, and SCS. This subset included 2,725 were from Ningxia and the remainder from Hebei.

The blood samples were used from the cows for extractions of the DNA using the DNeasy Blood & Tissue kit (QiaGen). We used 0.5 μg genomic DNA to build a paired end library according to the standard library preparation protocols, and sequencing was performed in DNBseq platform. The low-quality reads in 6010 cows were first filtered using fastp [25] with default parameters. Then, clean reads were mapped to the bovine reference genome ARS-UCD1.2 using BWA-MEM [26] v0.7.17. We used SAMtools [27] to convert and sort bam files.

### Data mapping

#### Feature engineering

In GP, the training population typically consists of several hundred individuals or samples, denoted as *n*, and millions of SNPs, denoted as *p*, with the condition *n* ≪ *p*. This imbalance poses a significant challenge for the performance of statistical models and machine learning-based methods [11]. One effective strategy to address this issue is reducing feature dimensionality by using annotated genes to represent the genome, thereby shifting from *p* to *q* (the number of annotated genes) [17]. However, the issue (*n* ≪ *q*) still persists. By considering each annotated gene as a haplotype, GP performance can be improved. Nevertheless, genes are typically longer than standard haplotypes, which might hinder performance. To mitigate this, gene regions can be divided into multiple haplotypes, allowing these haplotypes to represent the gene more effectively, thus enhancing GP performance.

To implement this approach, we utilized the ANNOVAR software [28] to annotate each gene with 16 variant types, including: ‘upstream’, ‘downstream’, ‘UTR’, ‘intronic’, ‘nonsynonymous’, ‘synonymous SNV’, ‘exonic’, ‘ncRNA exonic’, ‘ncRNA intronic’, ‘ncRNA splicing’, ‘stoploss’, ‘stopgain’, ‘splicing’. As well as two additional variant types extracted by GWAS: ‘gwas_postive’ and ‘gwas_negative’. A detailed description of these 16 variant types is provide in **Table S7**. The feature description for the *j*th gene in the *i*th sample (1 ≤ *i* ≤ *n*, 1 ≤ *j* ≤ *q*) is denoted as *g*_*ij*_, defined as:

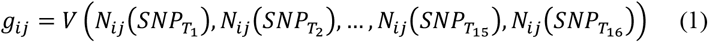

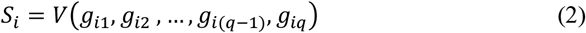

where 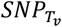 denotes the *v*th(*v* ∈ [1,16]) type of variant, and 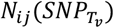 is the count of the *v*th (*v* ∈ [1,16]) type of variants in the *j*th gene for the *i*th sample. The vector function *V*(.) constructs a vector of 16 values representing 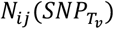. Consequently, *S*_*i*_ becomes a vector consisting of *q* vectors, forming a *q* × 16 matrix to describe the *i*th sample by genes instead of SNPs. This method achieves the first phase of dimensionality reduction from *p* to *q* in the dataset. The process is further detailed in **Algorithm 1**.

##### Algorithm 1. GENE-ENCODING(*x*)

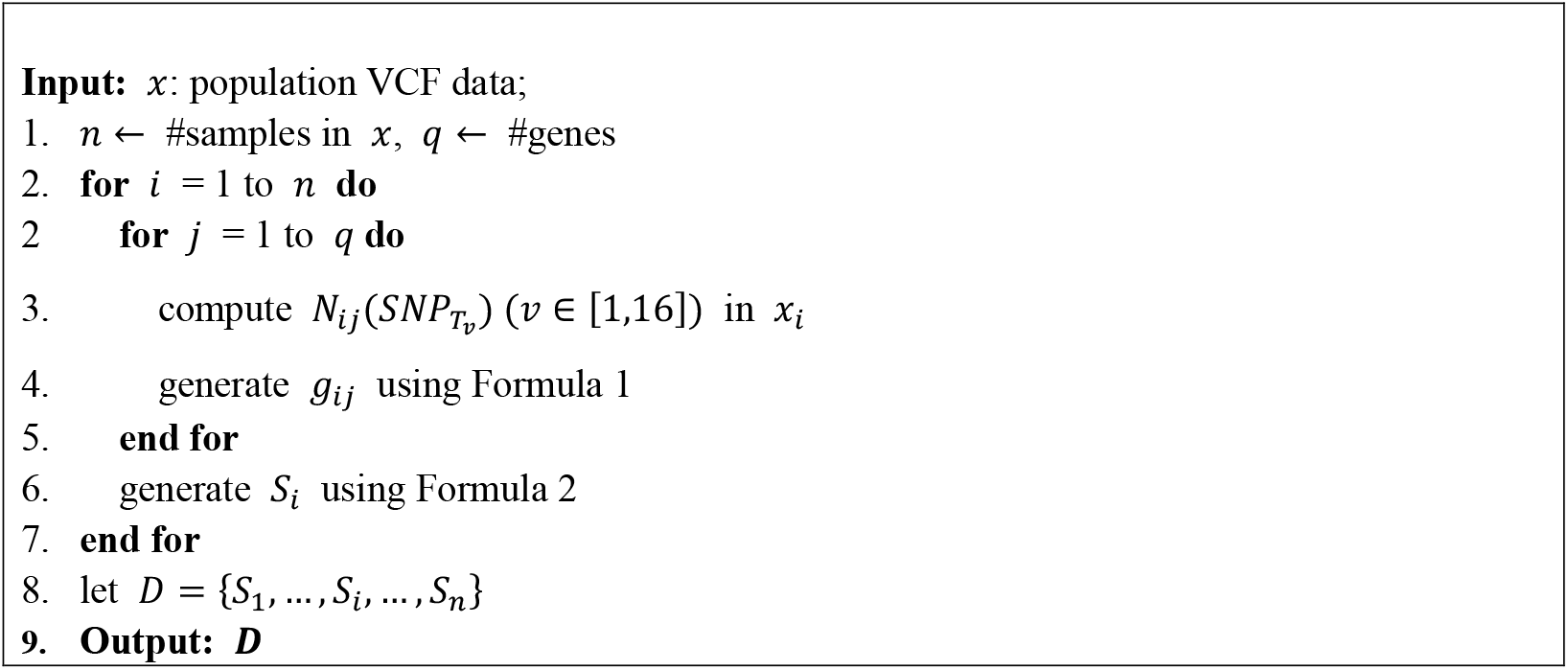

#### Data division

Based on **Algorithm 1**, the raw data of *n* samples is transformed into a three-dimensional data tensor, *D*(1.. *n*, 1.. *q*, 1. .16), defined as:

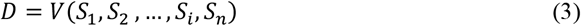

To effectively work with *D*, we developed a novel divide-and-conquer algorithm for GS, detailed in **Algorithm 2**. This algorithm consists of two phases: data division and data conquering. During the division phase (lines 4-5 in **Algorithm 2**), matrices *D*_*v*_(1.. *n*, 1.. *q, v*) are extracted for the *v* th feature (the *v* th type of variants) from *D*, where *v* ∈ [1,16]. Each *D*_*v*_(1.. *n*, 1.. *q, v*) denotes a *n* × *q* matrix, with each row represents a sample and each column corresponds to a gene, and the value in each cell indicating the occurrence count of the *v*th type of variants in the corresponding gene and sample. In this way, the three-dimensional data tensor *D* is divided into 16 two-dimensional matrices.

### Ensemble machine learning-based model

#### Dataset conquering

The prediction model in the proposed GP-ML-DC framework consists of two stages: conquering and combining, as detailed in **Algorithm 2**. In the conquering stage, we launch a subtask *M*_*v*_(1 ≤ *v* ≤ 16) by training the LASSO model on the extracted matrix *D*_*v*_(1.. *n*, 1.. *q, v*) for the *v*th type of feature (variant). Each *M*_*v*_ learns the contribution of the *v*th haplotype (e.g., some variant region, the *v* type of variants) of each gene to the phenotypes. The prediction outcome of *M*_*v*_ provides a partial contribution to the overall genome prediction task. By drawing on the principles of stacking ensemble learning [29], we regarded the prediction value of *M*_*v*_ as a meta-feature of the final genome prediction task. To mitigate overfitting, a ten-fold cross-validation strategy is employed to learn meta-features, the process (lines 6-15 in **Algorithm 2**) is described as follows. The matrix *D*_*v*_(1.. *n*, 1.. *q, v*) is randomly divided into ten folds (line 6 in **Algorithm 2**). A LASSO model is then trained on nine folds while predicting the remaining fold (lines 8-9 in **Algorithm 2**), with the prediction results serving as meta-features for the samples in the validation fold (lines 11-14 in **Algorithm 2**). This process repeats for all ten folds, resulting in ten trained models stored in *M*_*v*_ (line 16 in **Algorithm 2**), and converting *D*_*v*_(1.. *n*, 1.. *q, v*) into a one-dimensional vector of length *n*, where each element represents the meta-feature of a sample. Upon completing all 16 subtasks(*M*_1_, *M*_2_, …, *M*_16_), the three-dimensional dataset *D*(1.. *n*, 1.. *q*, 1. .16) is transformed into a new dataset where each sample is represented by a vector of 16 meta-feature values (line 13 in **Algorithm 2**). This process accomplishes the second phase of dimensionality reduction from *q* to 16 features.

##### Algorithm 2. ML-DIVIDE-CONQUER (*x*, y)

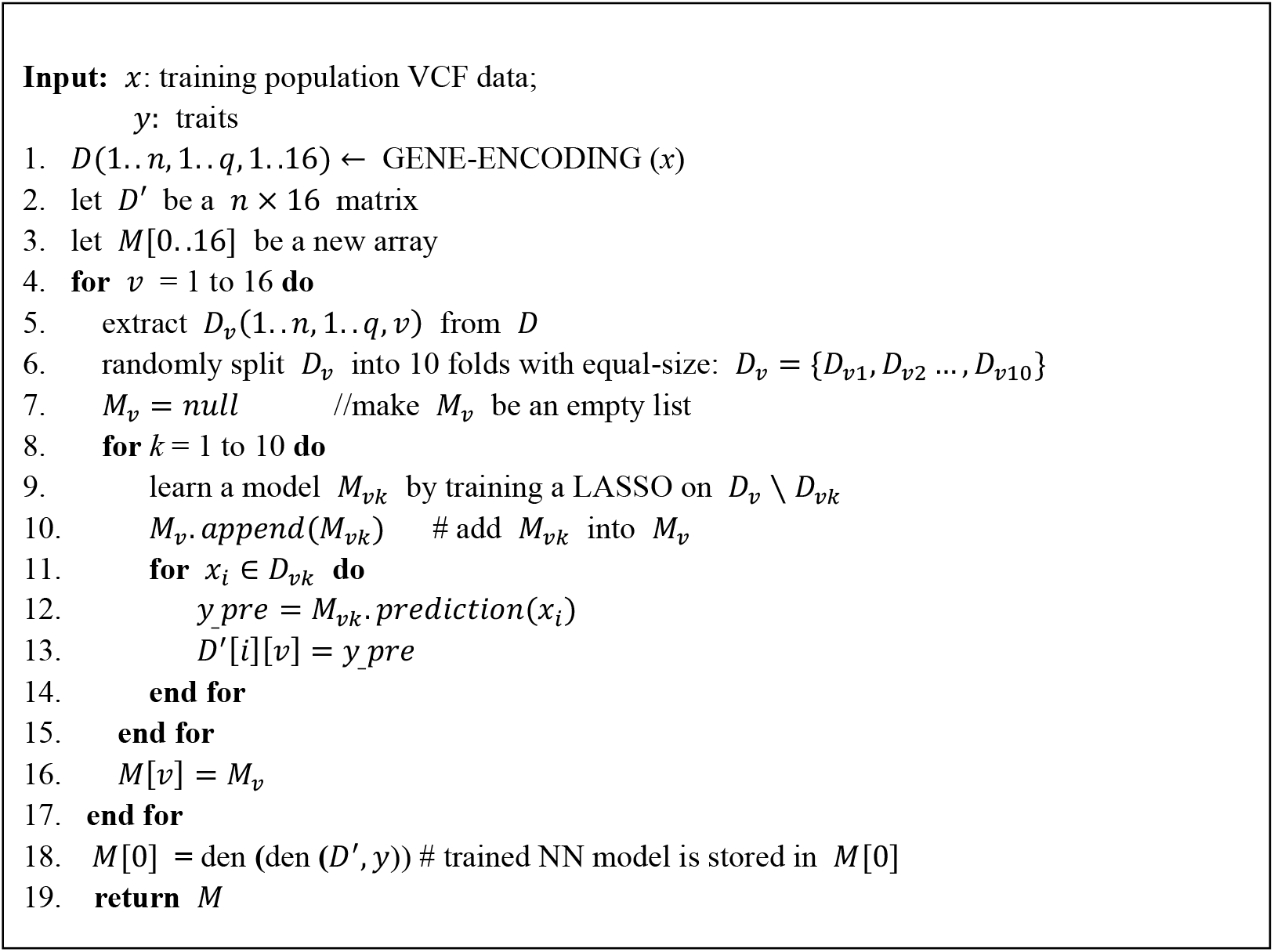

## Result combining

To integrate the contributions of all haplotypes (e.g., various types of variant regions) within genes, we trained a neural network with two fully connected layers on *D*′(1.. *n*, 1. .16). This network consolidates prediction results from each type of variant region (a haplotype) surrounding each gene for the final prediction. To enhance the representation capability of network. The first layer is followed by a ReLU activation function to enhance representational capacity. The output layer with a a single neuron generates the final prediction result. The process is described in line 17 of **Algorithm 2**.

### Prediction

For phenotype prediction of a testing population, given their variant calling genotype data *x_test*, and corresponding traits *y_train* from a training population *x_train* (variant calling genotype data) The phenotype prediction process for *x_test* is detailed in **Algorithm 3**. Initially, we apply the GENE-ENCODING (*x_test*) process on *x_test* as detailed in **Algorithm 1**, mapping *x_test* into *D*, which is characterized by genes. Following this, we invoke the ML**-**DIVIDE-CONQUER (*x_train,y_train*) method on *x_train* and *y_train* as outlined in **Algorithm 2**, producing a list of models, denoted as *M*. Here, *M*[0] (line 18 in the **Algorithm 2**) denotes the final model for genome prediction, while *M*[*v*](1 ≤ *v* ≤ 16) contains 10 models that are used to derive meta-features via ten-fold cross-validation for the *v*th type of variants (lines 7-10 in the **Algorithm 2**). Using *M*[*v*], predictions are made on *D*_*v*_ (the matrix denoting the *v*th type of feature in *D*) to generate meta-features (lines 10-18 in the **Algorithm 3**). The average of 10 prediction values provides the meta-feature’s value for each type of variants in testing dataset (line 19 in the **Algorithm 3**). Finally, *M*[0] is launched to predict the phenotype using the datasets consisting of meta-features as input (line 21 in the **Algorithm 3**).

### Extraction of variants from genes

Unlike the traditional GP methods that rely on SNP genotypes, the GP-ML-DC approach represents genetic information using genes as descriptors. However, SNPs embedded within these genes can still be extracted to form a subset. To evaluate the effectiveness of these extracted SNPs, we employ **Algorithm 4** to create a 50K SNP chip, allowing for comparison with the standard 50K chip typically used for analyzing milk traits in cows within the research community. In our analysis, a training population dataset *D*(1.. *n*, 1.. *q*, 1. .16), as defined in **Formula 3**, is organized using genes and can be derived using **Algorithm 1**. According to lines 3-8 in **Algorithm 4**, for the *v*th feature (the *v*th type of variants), where *v* ∈ [1,16], a matrix *D*_*v*_(1.. *n*, 1.. *q, v*) is extracted from *D*. A LASSO model is then trained on *D*_*v*_ to determine gene contributions to traits, resulting in a ranked list *l*_*v*_. Subsequently, in lines 10-17 in **Algorithm 4**, SNP variants from the top *k* genes in each *l*_*v*_(*v* ∈ [1,16]) are added to the *SNP*50 set, where *k* is less than the size of *l*_*v*_. SNPs in the *SNP*50 set are governed by a linkage disequilibrium (LD) threshold of 0.7, until the set *SNP*50 reaches or exceeds 50K.

#### Algorithm 3. GENOME-PREDICTION (*x_train, y_train, x_test*)

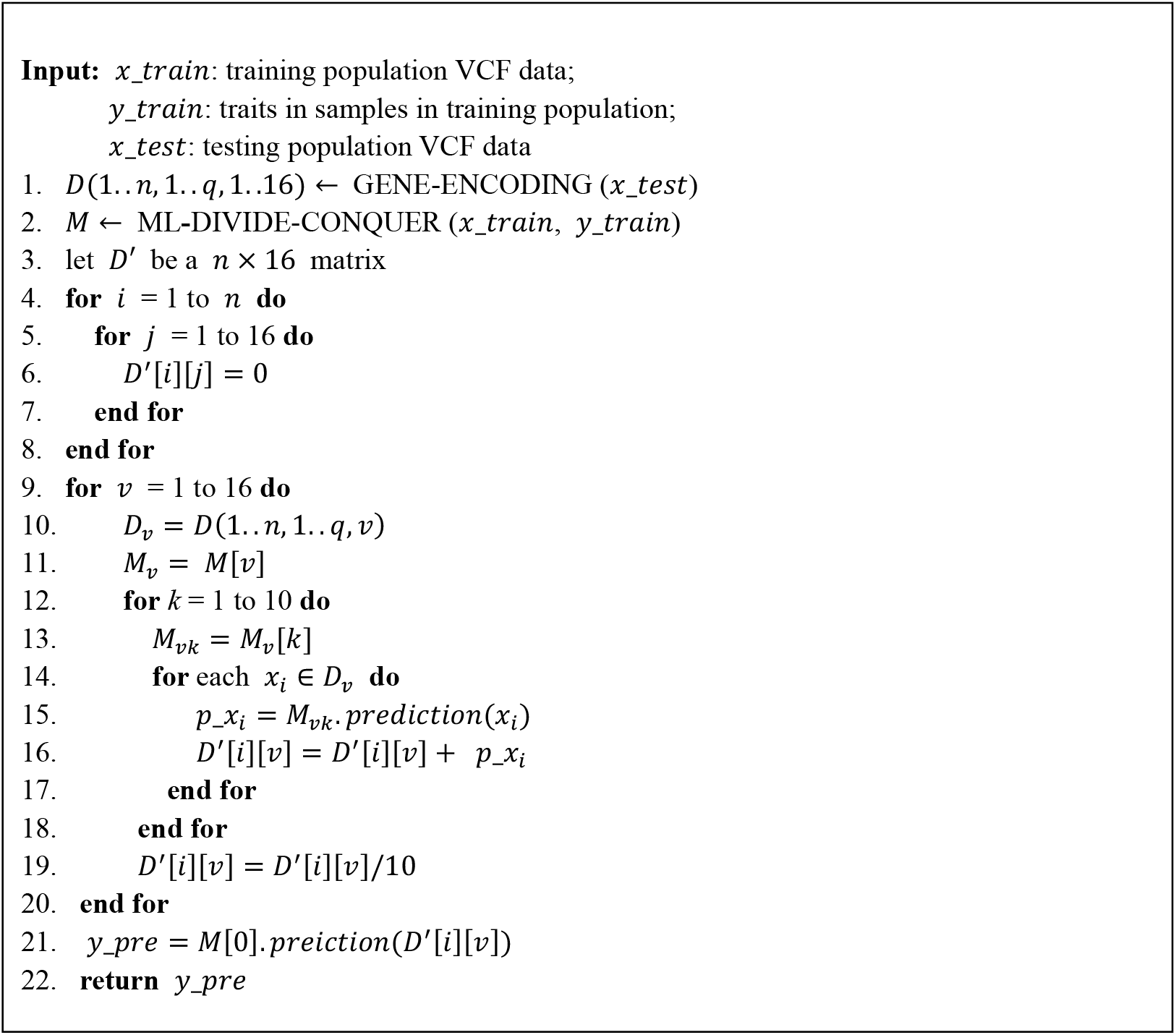

#### Algorithm 4. SNP50_Extraciton (*x*)

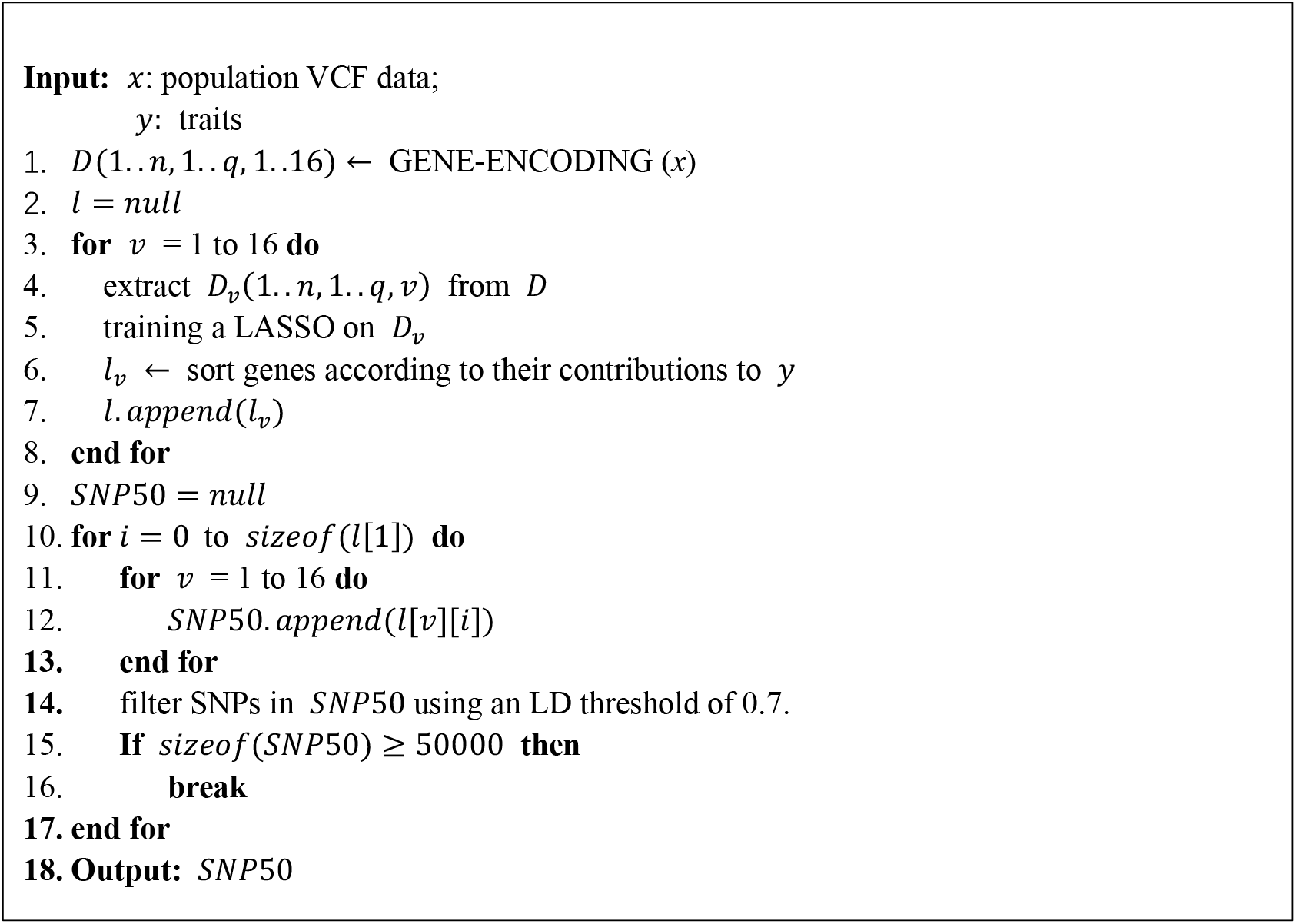

### Evaluation metrics and strategy

In the GS community, predictors typically fall into three categories: statistical models, ML-based models, and DL-based models. To evaluate the proposed model, we selected a state-of-the-art method from each category: the GBLUP statistics model, LightGBM [20] based on traditional ML, and DNNGP [21] based on DL. We employed two commonly used evaluation metrics in GS: Pearson’s correlation coefficient (PCC) and R-squared (R^2^), calculating these metrics to compare predicted and observed values in assessing model performance.

*k-*fold cross validation is a widely used strategy for evaluating GP models in the GS the research community. In this study, we conducted ten-fold cross validation on the Hebei dataset to benchmark the performance of GP-ML-DC against the three selected models. Each method underwent identical ten-fold partitioning. To mitigate potential bias from a single ten-fold cross validation, we performed five rounds of ten-fold cross-validation across four traits. The average results from these rounds were taken as the final performance measure.

It is essential to evaluate the generalization capability of ML models. While traditional ML and DL methods generally utilize SNPs-based genotype matrix for GP, inconsistent SNP spaces across individuals or groups can hinder their application in GP [17]. To further evaluate the effective of GP-ML-DC in predicting various individuals or populations, we conducted an independent test: predicting four traits in the Ningxia dataset using a model trained on the Hebei dataset.

## Acknowledgments

We thank the High-Performance Computing (HPC) of Northwest A&F University (NWAFU) for providing computing resource, as well as artificial Intelligence platform of the College of Information Engineering, Northwest A&F University.

