## Supplementary material for "GP-ML-DC: An Ensemble Machine Learning-Based Genomic Prediction Approach with Automated Two-Phase Dimensionality Reduction via Divide-and-Conquer Techniques": GP-ML-DC-Supp-7.17.16.30pm.docx

**Table S1**. Statistical difference of prediction performance (PCC) between GP-ML-DC and GBLUP on four traits for the CattleHeBei dataset.

| Traits | Variances | Levene's-Test for EV | | t-test for equality of mean | | | | | 95% Confidence Interval of the Difference | |
| --- | --- | --- | --- | --- | --- | --- | --- | --- | --- | --- |
|  |  | F | Sig. | t | df | Sig. (2-tailed) | md | sed | Lower | Upper |
| DMY | EVA | 20.079 | 2.0161E-6 | 13.929 | 98 | 5.7106E-25 | 0.099 | 0.007 | 0.084 | 0.112 |
|  | EVNA | - | - | 13.929 | 73.109 | 2.9151E-22 | 0.099 | 0.007 | 0.084 | 0.112 |
| MFY | EVA | 26.525 | 1.3448E-7 | 22.142 | 98 | 6.3764E-40 | 0.150 | 0.007 | 0.137 | 0.164 |
|  | EVNA | - | - | 22.142 | 68.986 | 4.5663E-33 | 0.150 | 0.007 | 0.137 | 0.164 |
| MPY | EVA | 17.878 | 5.2949E-6 | 12.261 | 98 | 1.6778E-21 | 0.093 | 0.008 | 0.078 | 0.108 |
|  | EVNA | - | - | 12.261 | 69.673 | 4.5958E-19 | 0.093 | 0.008 | 0.078 | 0.108 |
| SCS | EVA | 6.964 | 9.6746E-3 | 7.515 | 98 | 2.7014E-11 | 0.046 | 0.006 | 0.034 | 0.058 |
|  | EVNA | - | - | 7.515 | 87.405 | 4.6225E-11 | 0.046 | 0.006 | 0.034 | 0.058 |

Abbreviations: EV: Equality of Variances; EVA: Equal Variances Assumed; EVNA: Equal Variances Not Assumed; df: degrees of freedom; md: mean differences of PCC values of two models, a positive value shows the mean PCC of GP-ML-DC is higher than that of GBLUP; sed: standard error differences.

**Table S2**. Statistical difference of prediction performance (PCC) between GP-ML-DC and LightGBM on four traits for the CattleHeBei dataset.

| Traits | Variances | Levene's-Test for EV | | t-test for equality of mean | | | | | 95% Confidence Interval of the Difference | |
| --- | --- | --- | --- | --- | --- | --- | --- | --- | --- | --- |
|  |  | F | Sig. | t | df | Sig. (2-tailed) | md | sed | Lower | Upper |
| DMY | EVA | 16.537 | 9.648E-5 | 3.251 | 98 | 1.5747E-3 | 0.022 | 0.007 | 0.008 | 0.035 |
|  | EVNA | - | - | 3.251 | 76.281 | 1.7105E-3 | 0.022 | 0.007 | 0.008 | 0.035 |
| MFY | EVA | 12.588 | 5.979E-4 | 4.407 | 98 | 2.8619E-5 | 0.024 | 0.005 | 0.013 | 0.035 |
|  | EVNA | - | - | 4.407 | 81.501 | 3.1696E-5 | 0.024 | 0.005 | 0.013 | 0.035 |
| MPY | EVA | 11.831 | 8.5698E-4 | 0.424 | 98 | 0.672 | 0.003 | 0.006 | -0.009 | 0.014 |
|  | EVNA | - | - | 0.424 | 84.802 | 0.672 | 0.003 | 0.006 | -0.009 | 0.014 |
| SCS | EVA | 5.152 | 0.025 | -6.306 | 98 | 8.26E-9 | -0.039 | 0.006 | -0.051 | -0.026 |
|  | EVNA | - | - | -6.306 | 87.612 | 1.1278E-8 | -0.039 | 0.006 | -0.051 | -0.026 |

Abbreviations: EV: Equality of Variances; EVA: Equal Variances Assumed; EVNA: Equal Variances Not Assumed; df: degrees of freedom; md: mean differences of ACC values of two models, a positive value shows the mean PCC of GP-ML-DC is higher than that of LightGBM; sed: standard error differences.

**Table S3**. Statistical difference of prediction performance (PCC) between GP-ML-DC and DNNGP on four traits for the CattleHeBei dataset.

| Traits | Variances | Levene's-Test for EV | | t-test for equality of mean | | | | | 95% Confidence Interval of the Difference | |
| --- | --- | --- | --- | --- | --- | --- | --- | --- | --- | --- |
|  |  | F | Sig. | T | df | Sig. (2-tailed) | md | sed | Lower | Upper |
| DMY | EVA | 10.602 | 1.560E-3 | 15.262 | 98 | 1.2094E-27 | 0.104 | 0.007 | 0.09 | 0.118 |
|  | EVNA | - | - | 15.262 | 75.009 | 1.0429E-24 | 0.104 | 0.007 | 0.09 | 0.118 |
| MFY | EVA | 13.576 | 3.755E-4 | 18.868 | 98 | 2.1429E-34 | 0.122 | 0.006 | 0.109 | 0.135 |
|  | EVNA | - | - | 18.868 | 71.301 | 1.9458E-29 | 0.122 | 0.006 | 0.109 | 0.135 |
| MPY | EVA | 16.730 | 8.843E-5 | 19.523 | 98 | 1.5173E-35 | 0.132 | 0.007 | 0.119 | 0.145 |
|  | EVNA | - | - | 19.523 | 75.593 | 2.9741E-31 | 0.132 | 0.007 | 0.119 | 0.145 |
| SCS | EVA | 5.877 | 0.017 | 2.196 | 98 | 0.03 | 0.015 | 0.007 | 0.001 | 0.029 |
|  | EVNA | - | - | 2.196 | 79.445 | 0.031 | 0.015 | 0.007 | 0.001 | 0.029 |

Abbreviations: EV: Equality of Variances; EVA: Equal Variances Assumed; EVNA: Equal Variances Not Assumed; df: degrees of freedom; md: mean differences of ACC values of two models, a positive value shows the mean PCC of GP-ML-DC is higher than that of DNNGP; sed: standard error differences.

**Table S4**. Statistical difference of prediction performance (R^2^) between GP-ML-DC and GBLUP on four traits for the CattleHeBei dataset.

| Traits | Variances | Levene's-Test for EV | | t-test for equality of mean | | | | | 95% Confidence Interval of the Difference | |
| --- | --- | --- | --- | --- | --- | --- | --- | --- | --- | --- |
|  |  | F | Sig. | t | df | Sig. (2-tailed) | md | sed | Lower | Upper |
| DMY | EVA | 11.498 | 0.001 | 14.522 | 98 | 3.5961E-26 | 0.107 | 0.007 | 0.092 | 0.121 |
|  | EVNA | - | - | 14.522 | 81.466 | 2.57E-24 | 0.107 | 0.007 | 0.092 | 0.121 |
| MFY | EVA | 8.441 | 0.005 | 25.171 | 98 | 1.4352E-44 | 0.147 | 0.006 | 0.136 | 0.159 |
|  | EVNA | - | - | 25.171 | 85.622 | 2.4677E-41 | 0.147 | 0.006 | 0.136 | 0.159 |
| MPY | EVA | 10.287 | 0.002 | 12.974 | 98 | 5.3275E-23 | 0.098 | 0.008 | 0.083 | 0.114 |
|  | EVNA | - | - | 12.974 | 78.264 | 3.3867E-21 | 0.098 | 0.008 | 0.083 | 0.114 |
| SCS | EVA | 0.461 | 0.499 | 7.462 | 98 | 3.5013E-11 | 0.051 | 0.007 | 0.037 | 0.064 |
|  | EVNA | - | - | 7.462 | 76.017 | 1.1707E-10 | 0.051 | 0.007 | 0.037 | 0.064 |

Abbreviations: EV: Equality of Variances; EVA: Equal Variances Assumed; EVNA: Equal Variances Not Assumed; df: degrees of freedom; md: mean differences of ACC values of two models, a positive value shows the mean R^2^ of GP-ML-DC is higher than that of GBLUP; sed: standard error differences.

**Table S5**. Statistical difference of prediction performance (R^2^) between GP-ML-DC and LightGBM on four traits for the CattleHeBei dataset.

| Traits | Variances | Levene's-Test for EV | | t-test for equality of mean | | | | | 95% Confidence Interval of the Difference | |
| --- | --- | --- | --- | --- | --- | --- | --- | --- | --- | --- |
|  |  | F | Sig. | t | df | Sig. (2-tailed) | md | sed | Lower | Upper |
| DMY | EVA | 14.997 | 1.9444E-4 | 3.295 | 98 | 1.3704E-3 | 0.026 | 0.008 | 0.01 | 0.041 |
|  | EVNA | - | - | 3.295 | 78.027 | 1.4816E-3 | 0.026 | 0.008 | 0.01 | 0.041 |
| MFY | EVA | 9.594 | 0.003 | 4.356 | 98 | 3.2584E-5 | 0.025 | 0.006 | 0.014 | 0.037 |
|  | EVNA | - | - | 4.356 | 85.342 | 3.6641E-5 | 0.025 | 0.006 | 0.014 | 0.037 |
| MPY | EVA | 10.449 | 0.002 | 0.689 | 98 | 0.4925 | 0.005 | 0.007 | -0.009 | 0.018 |
|  | EVNA | - | - | 0.689 | 86.417 | 0.4927 | 0.005 | 0.007 | -0.009 | 0.018 |
| SCS | EVA | 0.441 | 0.508 | -1.646 | 98 | 0.1029 | -0.015 | 0.009 | -0.034 | 0.003 |
|  | EVNA | - | - | -1.646 | 94.338 | 0.1029 | -0.015 | 0.009 | -0.034 | 0.003 |

Abbreviations: EV: Equality of Variances; EVA: Equal Variances Assumed; EVNA: Equal Variances Not Assumed; df: degrees of freedom; md: mean differences of ACC values of two models, a positive value shows the mean R2 of GP-ML-DC is higher than that of LightGBM; sed: standard error differences.

**Table S6**. Statistical difference of prediction performance (R^2^) between GP-ML-DC and DNNGP on four traits for the CattleHeBei dataset.

| Traits | Variances | Levene's-Test for EV | | t-test for equality of mean | | | | | 95% Confidence Interval of the Difference | |
| --- | --- | --- | --- | --- | --- | --- | --- | --- | --- | --- |
|  |  | F | Sig. | t | df | Sig. (2-tailed) | md | sed | Lower | Upper |
| DMY | EVA | 5.016 | 0.027 | 16.086 | 98 | 2.9977E-29 | 0.113 | 0.007 | 0.098 | 0.126 |
|  | EVNA | - | - | 16.086 | 84.825 | 1.6987E-27 | 0.113 | 0.007 | 0.098 | 0.126 |
| MFY | EVA | 5.951 | 0.017 | 20.264 | 98 | 8.0745E-37 | 0.123 | 0.006 | 0.111 | 0.135 |
|  | EVNA | - | - | 20.264 | 83.281 | 6.1259E-34 | 0.123 | 0.006 | 0.111 | 0.135 |
| MPY | EVA | 6.777 | 0.011 | 20.056 | 98 | 1.8241E-36 | 0.133 | 0.007 | 0.119 | 0.145 |
|  | EVNA | - | - | 20.056 | 87.918 | 1.4621E-34 | 0.133 | 0.007 | 0.119 | 0.145 |
| SCS | EVA | 0.725 | 0.397 | 3.886 | 98 | 1.8510E-4 | 0.030 | 0.008 | 0.015 | 0.045 |
|  | EVNA | - | - | 3.886 | 94.014 | 1.8957E-4 | 0.030 | 0.008 | 0.015 | 0.045 |

Abbreviations: EV: Equality of Variances; EVA: Equal Variances Assumed; EVNA: Equal Variances Not Assumed; df: degrees of freedom; md: mean differences of ACC values of two models, a positive value shows the mean R^2^ of GP-ML-DC is higher than that of DNNGP; sed: standard error differences.

**Table S7.** 16 types of variant features for gene description

|  | Type | Descript |
| --- | --- | --- |
| 1 | upstream | The variant spans a 1-kb region upstream of the transcription start site |
| 2 | downstream | The variant overlaps a 1-kb region downstream of the transcription end site |
| 3 | UTR | The variant overlaps a 5' untranslated region and a 3' untranslated region |
| 4 | intronic | The variant overlaps an intron |
| 5 | nonsynonymous | A single nucleotide change that results in an amino acid alteration |
| 6 | synonymous SNV | A single nucleotide change that results in no amino acid alteration |
| 7 | exonic | The variant overlaps with a coding region |
| 8 | ncRNA exonic | The variant overlaps with a noncoding exonic region |
| 9 | ncRNA intronic | The variant overlaps with a noncoding intronic region |
| 10 | ncRNA splicing | The variant overlaps with a noncoding splicing region |
| 11 | stoploss | A nonsynonymous SNV, frameshift insertion/deletion, nonframeshift insertion/deletion, or block substitution that results in the immediate removal of a stop codon at the variant site. |
| 12 | stopgain | A nonsynonymous SNV, frameshift insertion/deletion, nonframeshift insertion/deletion, or block substitution that results in the immediate emergence of a stop codon at the variant site. In the case of frameshift mutations, the appearance of a stop codon downstream of the variant will not be classified as "stopgain." |
| 13 | splicing | The variant is within 2 base pairs of a splicing junction |
| 14 | eQTL | The expression quantitative trait loci |
| 15 | gwas_postive | The variant has a positive effect on GWAS |
| 16 | gwas_negative | The variant has a negative effect on GWAS |
